# A linear-time algorithm to sample the dual-birth model

**DOI:** 10.1101/226423

**Authors:** Niema Moshiri

## Abstract

The ability to sample models of tree evolution is essential in the analysis and interpretation of phylogenetic trees. The dual-birth model is an extension of the traditional birth-only model and allows for sampling trees of varying degrees of balance. However, for a tree with *n* leaves, the tree sampling algorithm proposed in the original paper is 𝒪(*n* log *n*). I propose an algorithm to sample trees under the dual-birth model in 𝒪(*n*), and I provide a fast C++ implementation of the proposed algorithm.

## I INTRODUCTION

The dual-birth model is a birth-only model in which each branch is a Poisson process with one of two states: *active,* in which splits occur with rate λ_*b*_, and *inactive,* in which splits occur with rate λ_*a*_. It is typically parameterized by λ = λ_*a*_ + λ_*b*_ and *r* = λ_*a*_/λ_*b*_ [1]. When a branch splits, one child branch is active and the other is inactive. Simulating trees under the dual-birth model can provide null distributions describing neutral evolutionary process, which can be rejected when inferring trees from the data. Further, Maximum Likelihood (ML) tree inference techniques tend to infer significantly overbalanced trees when true trees are extremely unbalanced [1], and the dual-birth model allows for researchers to simulate trees of varying balance by adjusting its *r* parameter, making the dual-birth model useful for simulation experiments that study ML tree inference techniques. The algorithm originally proposed is 𝒪(*n* log *n*), where *n* is the number of leaves of the tree, but by exploiting statistical properties of Poisson processes and the exponential distribution, the sampling problem can be solved in 𝒪(*n*).

## II. METHODS

### A Linear-time algorithm description

Let exp (λ) denote an exponential random variable with rate λ. Let bern(*p*) denote an exponential random variable with probability *p*. Let ||*x*|| denote the size of set *x*. The algorithm starts with two sets of leaves, *active* (with just the root) and *inactive* (initially empty). The algorithm iterates while the termination condition has not yet been met. The time until the next split is sampled from exp (λ_*b*_||*active*|| + λ_*a*_||*inactive*||). The next splitting leaf is chosen by first determining its state by sampling bern (λ_*b*_||*active*||(λ_*b*_||*active*|| + λ_*a*_||*inactive*||)) (success = active, failure = inactive) and then uniformly choosing a random leaf from the corresponding set. The chosen leaf's time is set to the sampled split time, the leaf is removed from its set, two children are created, and one child is placed in *active* and the other in *inactive*. When the termination condition has been met, the times of all remaining leaves in *active* and *inactive* are set to the end time.

### B Algorithm correctness

The algorithm begins with a single active leaf at time 0. Because each branch is a Poisson process, the time until the branch splits is an exponential random variable with the branch’s rate. Therefore, at a given time *t* with *n_a_* active leaves and *n_i_* inactive leaves, the time until the next splitting event is the minimum of *n*_a_ exponential random variables, each with rate λ_*b*_, and *n_i_* exponential random variables, each with rate λ_*a*_.

The minimum of *k* exponential random variables *X*_1_,…, *X_k_* with rates λ_1_,…, λ_*k*_ is an exponential random variable with rate 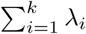. Therefore, the time until the next splitting event is an exponential random variable with rate *n*_*a*_λ_*b*_ + *n*_i_λ_*a*_.

Given *k* exponential random variables *X_1_,…, X_k_* with rates λ_1_,…,λ_*k*_, the probability that *X_i_* is the minimum of all *k* random variables is 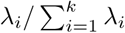. Because only one terminal branch can be the next to split, the events that pendant branch *j* (for all *j*) is the next to split are mutually exclusive. Therefore, the probability that the next branch to split is an active branch is *p*= *n*_*a*_λ_*b*_/(*n_a_*λ_*b*_+*n_i_*λ_*a*_). Once the state of the next branch to split is determined (by sampling a Bernoulli random variable with probability *p*), because all leaves in that state have the same rate (and are thus equally likely to be the next to split), the next splitting leaf can be chosen uniformly from the set of leaves in the chosen state.

## III. RESULTS

My implementation of the proposed algorithm closely follows the theoretical expectations of the dual-birth model (Fig. 1) and is orders of magnitude faster than the original implementation of the initially-proposed algorithm (Fig. 2).

**Fig. 1.**
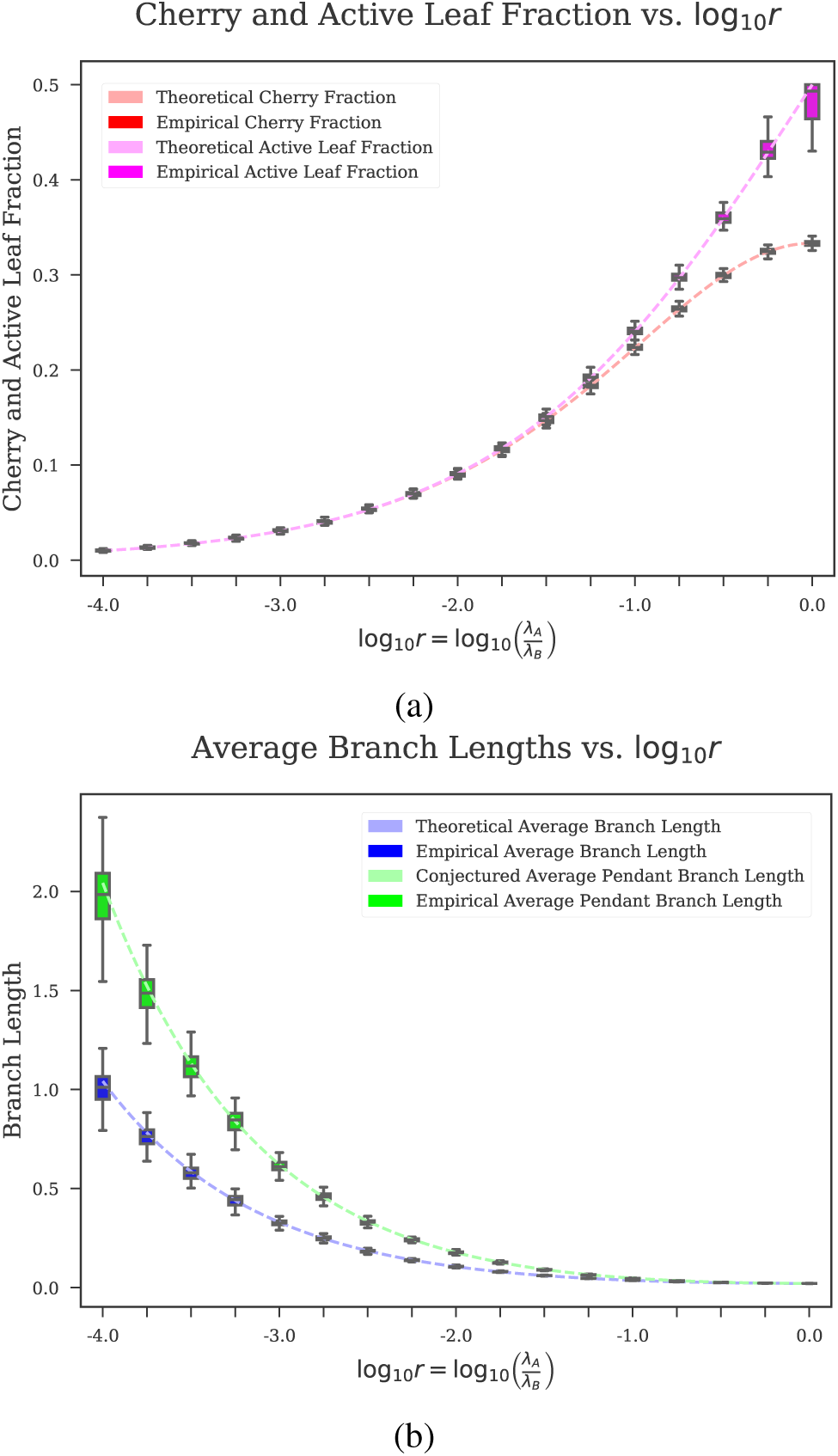
Theoretical expectations of (a) cherry fraction (dashed red line) and active leaf fraction (dashed purple line), and (b) branch length (dashed blue line) and terminal branch length (dashed green line) versus simulated distributions (in box plots) using 100 replicates with *n* = 4096, λ = 48, and varying values of *r* (*x*-axis) from 1/1024 to 1.

**Fig. 2.**
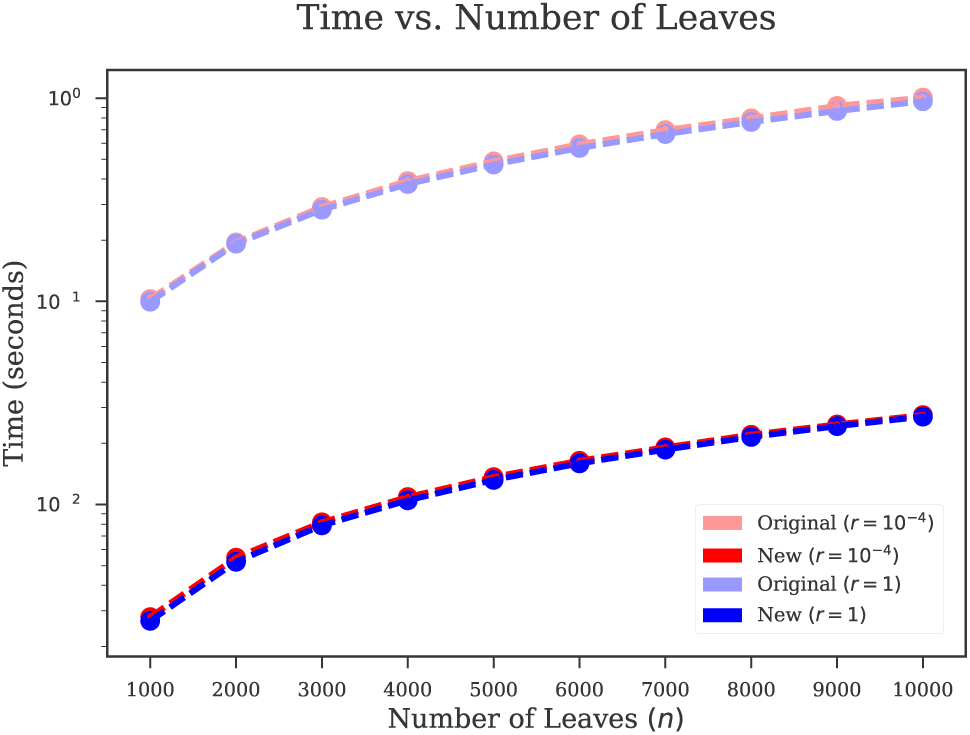
Log-scaled runtimes of the original Python-implemented algorithm and the new C++-implemented algorithm for 1000 ≤ *n* ≤ 10000 for *r* = 10^−4^ and *r* =1. Each measured time is the average of the time taken to simulate 20 trees in a single execution, and each point plot is the distribution of 20 replicate executions.

## IV. DISCUSSION

The proposed algorithm scales linearly with the number of leaves, and its C++ implementation is fast and does not have any non-STL dependencies, making it easy to build on all environments. My hope is that the fast implementation I provide can be wrapped into existing tools for the convenience of users as well as developers.

## AVAILABILITY

My C++ implementation of the proposed algorithm can be found in the following GitHub repository: https://github.com/niemasd/Dual-Birth-Simulator

## ACKNOWLEDGMENT

I thank Siavash Mirarab for his guidance in the original dual-birth model project.

